# Monitoring for SARS-CoV-2 drug resistance mutations in broad viral populations

**DOI:** 10.1101/2022.05.27.493798

**Authors:** Mayya Sedova, Lukasz Jaroszewski, Mallika Iyer, Adam Godzik

## Abstract

The search for drugs against COVID-19 and other diseases caused by coronaviruses focuses on the most conserved and essential proteins, mainly the main (M^pro^) and the papain-like (PL^pro^) proteases and the RNA-dependent RNA polymerase (RdRp). Nirmatrelvir, an inhibitor for M^pro^, was recently approved by FDA as a part of a two-drug combination, Paxlovid, and many more drugs are in various stages of development. Multiple candidates for the PL^pro^ inhibitors are being studied, but none have yet progressed to clinical trials. Several repurposed inhibitors of RdRp are already in use. We can expect that once anti-COVID-19 drugs become widely used, resistant variants of SARS-CoV-2 will emerge, and we already see that for the drugs targeting SARS-CoV-2 RdRp. We hypothesize that emergence of such variants can be anticipated by identifying possible escape mutations already present in the existing populations of viruses. Our group previously developed the coronavirus3D server (https://coronavirus3d.org), tracking the evolution of SARS-CoV-2 in the context of the three-dimensional structures of its proteins. Here we introduce dedicated pages tracking the emergence of potential drug resistant mutations to M^pro^ and PL^pro^, showing that such mutations are already circulating in the SARS-CoV-2 viral population. With regular updates, the drug resistance tracker provides an easy way to monitor and potentially predict the emergence of drug resistance-conferring mutations in the SARS-CoV-2 virus.

## Introduction

In the first two years of the COVID-19 pandemic, most efforts to combat its spread have focused on behavioral-based prevention measures (mask mandates, quarantines) and vaccine development. Once vaccines became available, the focus shifted to vaccination strategies and protocols. However, in the long-term, we also need to develop antiviral drugs, ideally with broad specificity, targeting all SARS-CoV-2 variants and possibly other related coronaviruses. An extensive research effort to find such drugs started almost immediately after the onset of the COVID-19 pandemic. After many false starts and bogus claims [2], the first targeted COVID-19 antivirals are now being approved and are entering clinical practice [3].

Borrowing from strategies used to combat other viruses, most of the efforts in drug discovery against COVID-19 focus on SARS-CoV-2 proteases (M^pro^ and PL^pro^) and RNA polymerase (RdRp). Broad spectrum antivirals targeting the latter, remdesivir and molnupiravir, were given emergency use authorization (EUA) in November 2020 and November 2021. They were initially developed as drugs against hepatitis C and flu, and both have significant side effects limiting their broader use. The first approved M^pro^ inhibitor developed specifically against COVID-19 by Pfizer is nirmatrelvir (PF-07321332, Bexovid) [4]. It received FDA emergency usage authorization (EUA) in December 2021 after a successful clinical trial [5] and, in combination with another protease inhibitor, ritonavir [6], is now available under the brand name Paxlovid. No PLpro inhibitors have been approved as of this writing; however, several of them, such as ebselen [7], are now in advanced clinical trials.

Like for every drug, we can expect the eventual emergence of drug resistance, brought about by mutations selectively blocking the effects of the drug. Growing resistance to common antibacterial drugs is one of the main medical challenges of the 21^st^ century [8]. While less talked about, the same problem emerged in all existing antiviral therapies. The emergence of resistance against antiviral treatments has been documented in the hepatitis B virus, human immunodeficiency virus (HIV-1), hepatitis C virus (HCV), and influenza virus [4]. Drug resistance mutations against remdesivir have already been identified in SARS-CoV-2 in *in vitro* experiments [9] and individual patients [10].

In most cases, drug resistance mutations are discovered as a response to the wide use of a drug or in laboratory escape studies when selection pressure leads to the rapid growth of resistant strains. However, genetic variations, especially recurrent ones, at the positions involved in drug binding in the bacterial or viral population prior to the drug release may give us a preview of the future development of drug resistance. The existence of such mutations in the drug-naïve population suggests that the pathogen tolerates modifications at these respective positions. Thus they can be quickly “rediscovered” by the virus once the selective pressure from the drug is applied.

## Results

Our group previously developed the Coronavirus3D server (https://coronavirus3d.org/) to analyze the distribution of SARS-CoV-2 mutations in the context of experimental or predicted 3D structures of its proteins [11]. It was later expanded to provide tracking of the dynamics of the emergence of novel variants and mutations as a part of the SARS-CoV-2 Assessment of Viral Evolution (SAVE) program [12]. SARS-CoV-2 genomes are downloaded regularly from the GISAID database [13], and the distribution and time patterns of mutations are analyzed to identify fast-growing or track otherwise important mutations. The server’s default page focuses on identifying the fastest growing SARS-CoV-2 variants, with information on genome modifications and their combinations available from the pull-down menus. The original pages on the 3D distribution of mutations [11] are available from the “3D proteome viewer” tab, while here, we describe newly developed pages (“Drug resistance” tab) providing information on the mutations that could potentially affect the efficacy of drugs targeting SARS-CoV-2 virus proteins. We initially focus on the M^Pro^, as a major drug target of an already approved drug with a well-established inhibition mechanism. In principle, any other proteins for which structures with the drug are available could be analyzed in the same way. Selected PL^pro^ inhibitors, currently in early research stages, are also included.

SARS coronavirus main protease (M^pro^) or 3C-like protease (3CL^pro^), also referred to as nonstructural protein 5 (nsp5), cleaves the coronavirus polyprotein into individual functional proteins, and its inhibition would stop the virus proliferation. It is a chymotrypsin-like cysteine protease, a member of the MEROPS [14] peptidase family C30 (clan PA(C)), and its active site consists of a Cys-His catalytic dyad (H41 and C145) and several binding pockets defining its cleavage specificity [15]. Homologous proteases with similar functions are found in the most positive sense, single-stranded RNA viruses ((+)ssRNA) viruses [16], and more distant homologs are found in organisms from all kingdoms of life [14]. A high level of active site conservation among coronaviruses opened the possibility of developing a broad specificity inhibitor targeting all coronaviruses, including other pathogenic human coronaviruses [4].

With over 10M SARS-CoV-2 genomes sequences available in the GISAID database [13] we have a deep insight into the potential mutability of each position in the genome. As shown by us [17] and others [18], nonstructural proteins of SARS-CoV-2 are under strong negative selection. As a result, they have significantly fewer mutations in relation to their length than structural proteins, especially spike glycoprotein. Nsp5 is one of the least often mutated proteins in the SARS-CoV-2, consistent with its essential role in the virus’s lifecycle. Still, many positions in M^pro^ have been mutated and some of these mutations, such as P132H, are founders of some variants of concern, here the Omicron variant. However, this mutation is far away from the active/binding site and does not seem to offer any advantage to the virus [19].

The dynamics of the COVID-19 pandemic have been dominated by the subsequent waves of novel variants, the largest of which are called variants of concern (VOC), and were given easy-to-remember names based on a Greek alphabet. Focus on the several named VOSs underplays the genomic diversity of the SARS-CoV-2, with over two thousand named variants [20] and many recurrent mutations, deletions, and insertions concentrated in several highly variable regions [21], but also present throughout the genome. Among the mutations observed in the SARS-CoV-2 genomes sequenced from patient samples, many may not offer any advantage to the virus, but all represent viable viruses. Mirroring the situation in cancer [22], such “passenger mutations” are tolerated as long as they are only mildly deleterious, adding to the genomic diversity of the population. When the evolutionary pressure changes, like after introducing a new drug, previously neutral or mildly deleterious mutation may provide an advantage to the virus. Therefore, the presence of mutations in the drug binding sites in the drug naïve viral population, still not under selection, may hint at the existence of easy escape routes for the virus. With massive tracking of the COVID-19 epidemics by genomic sequencing, we have unprecedented insight into the presence of such mutations.

The coronavirus3D server tracks potential resistance mutations by first identifying residues with any heavy atom within a specific distance from any atom of the inhibitor molecule. Several experimental structures of SARS-CoV-2 M^pro^ with nirmatrelvir [4] are available in the PDB database [23], one of which (7vh8 [24]) is used here as an example. We hypothesize that any mutation in proximity to the bound inhibitor may potentially affect the binding energy and, thus, the drug efficacy. Therefore, we use the regularly updated list of SARS-CoV-2 genomes to survey the presence and dynamics of the potential drug escape mutations in the SARS-CoV-2 genomes.

Figure 1 shows the main section of the drug resistance tab on the coronavirus3D server. The 3Dmol.js [25] visualization of the M^pro^ structure with the inhibitor is shown on the left. The ribbon diagram of the structure is colored by the density of mutations according to the color scheme that could be selected from the corresponding menus on the right. Other menus allow the user to specify the type of visualization, color schemes, select a specific set of PDB coordinates (currently from the fixed list), and the specific viral variant of concern to be analyzed. The default page on the sever focuses on the M^pro^ and its interactions with nirmatrelvir. Several entries with various drug candidates targeting PL^pro^ are also available.

**Figure 1.**
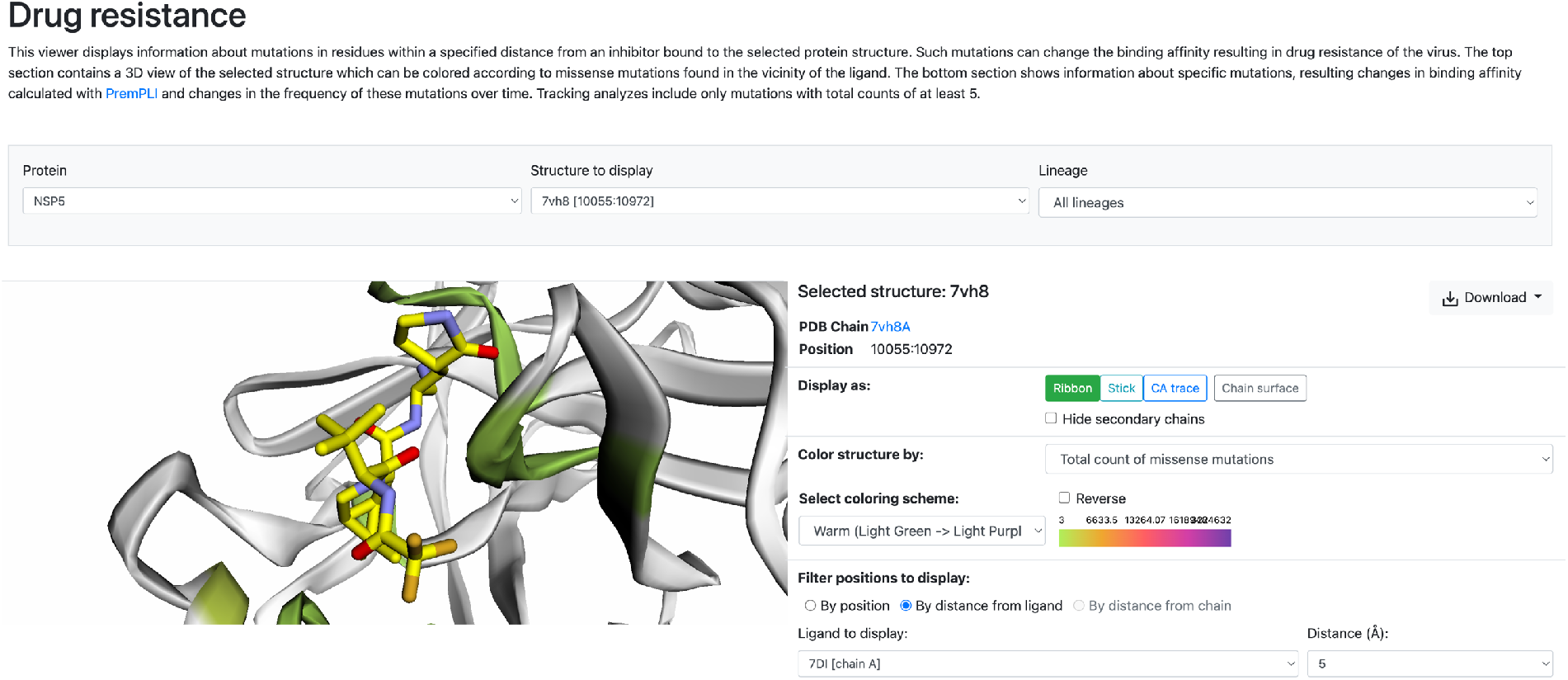
Coronavirus3D drug resistance page at https://coronavirus3d.org/#/drug, showing a picture of SARS-CoV-2 M^pro^ with a bound nirmatrelvir. M^pro^ structure is colored by the frequency of mutations in the regions within 5Å from any of the heavy atoms of the inhibitor.

Figure 2 shows the detailed report about individual mutations that includes tracking the absolute number of genomes with the given mutation and growth trends among such genomes collected in the last two years (in the “Recent growth” column). The histogram is colored based on the estimated effects of the mutations on the ligand here, the nirmatrelvir, provided by the PremPLI algorithm [26], a machine learning model for predicting the effects of missense mutations on protein-ligand interactions. The histogram, genome counts, and growth/decline trends, as well as other pages and tabs on the coronavirus3D server are updated regularly with the updates in the GISAID. The figures and counts presented here are based on the update from May 20^th^.

**Figure 2.**
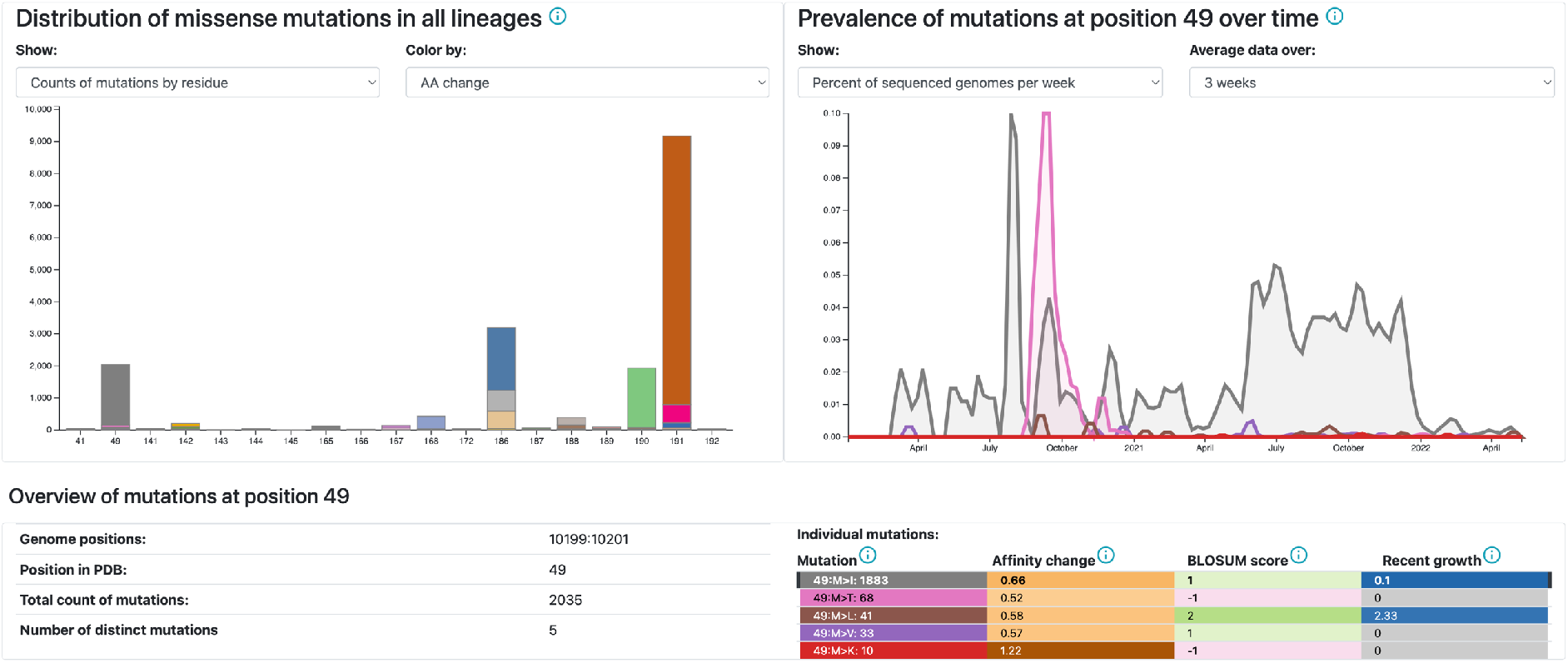
Tracking data on the mutations in the positions within a specified radius around the nirmatrelvir molecule (here 5Å, as selected on the top of the page (Figure 1)), based on the PDB structure 7vh8. Mutation counts are shown as histograms, and detailed information about individual mutations, including estimates of binding energy changes and time patterns of incidence becomes available after hovering over the respective element in the histogram. An example of a relatively common (1883 cases in GIDAID) M49I mutation is shown in Figure 3

**Figure 3.**
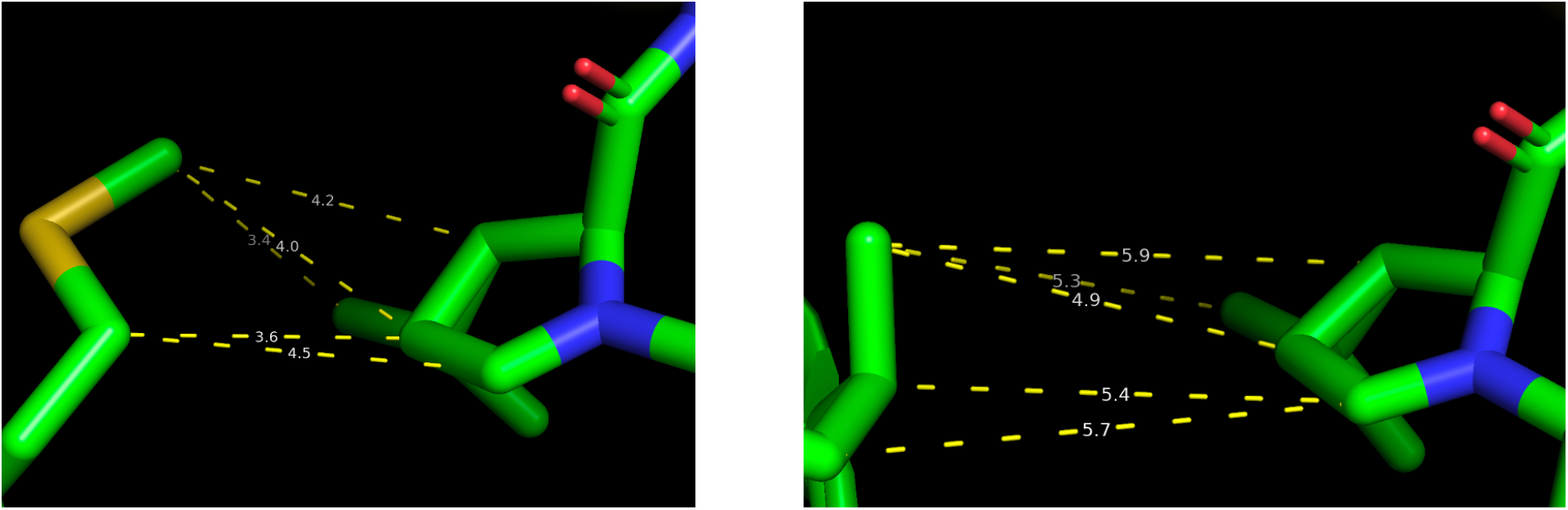
Interactions between residue M49 in the experimental structure of M^pro^ with nirmatrelvir (left panel) and the same interaction in the EvoEF [1] model of the M^pro^ with the M49I mutation. The M49I mutation was found in 1883 genomes, with a slight uptick in late 2021 (see Figure 2). The model predicts that mutation from methionine to smaller isoleucine results in a loss of close interactions. The ΔΔG of this mutation was predicted to be +0.66 kcal/mol.

As seen on Figures 1 and 2, mutations on the positions in the M^pro^ active site cavity are relatively rare, but they still include hundreds or even thousands of cases. It is worth noting that while mutations of the catalytic dyad are extremely rare (2 cases each, both in low-quality genomes, so likely just sequencing errors), positions on the edges of the active site cavity display a significant degree of mutability with mutations that are predicted to affect binding affinity to nirmatrelvir and thus its efficacy as a drug. In relatively small numbers, they were found across multiple variants and at different times, consistent with the picture of random fluctuations with no evolutionary pressure to be fixed. On the other hand, their presence in hundreds or thousands of genomes suggests that they are not particularly detrimental, so with the presence of the drug, they could expand with minimal cost to the virus. It is also worth noting that, while the absolute numbers are small, some of these potential escape mutations seem to be trending up in some of the recent Omicron subvariants.

## Conclusions

The unprecedented level of the genetic surveillance of the SARS-CoV-2 virus provided us with a wealth of information on the patterns and dynamics of mutations and other genomic modifications in the SARS-CoV-2 virus. Most of the attention focused on mutation signatures of the novel, major variants of SARS-CoV-2, and their ability to avoid immune memory from vaccinations or previous infections and other regions under positive selection. Here we argue that regions and mutations under neutral or weak negative selection may provide a hint about the future behavior of the virus, especially if we have reasons to expect that some of such mutations may become beneficial for the virus in some circumstances. The release of new anti-COVID-19 treatments can create such circumstances when the virus facing pressure from a drug could rapidly increase subpopulations with the existing or recurring escape mutations. With recent releases of new antiviral treatments such as Paxlovid and in anticipation of other releases, we developed a series of tracking pages where positions involved in interactions with the drugs are monitored for (currently) neutral mutations. This information can be used to predict future possible escape mechanisms of SARS-CoV-2 and to design the next generation of anti-COVID-19 drugs.

## Acknowledgments

We gratefully acknowledge the Authors from the Originating laboratories responsible for obtaining the specimens and the submitting laboratories where genetic sequence data were generated and shared via the GISAID Initiative, on which this research is based. We further acknowledge comments and suggestions from our colleagues from the CSGID collaboration and the SAVE program. This research was partly supported by NIH-NIAID Center for Structural Genomics of Infectious Diseases (CGGID) contract award HHSN272201700060C.

